## Appendix 1 for "Structural basis for molecular assembly of fucoxanthin chlorophyll *a*/*c*-binding proteins in a diatom photosystem I supercomplex"

### Supplementary Information

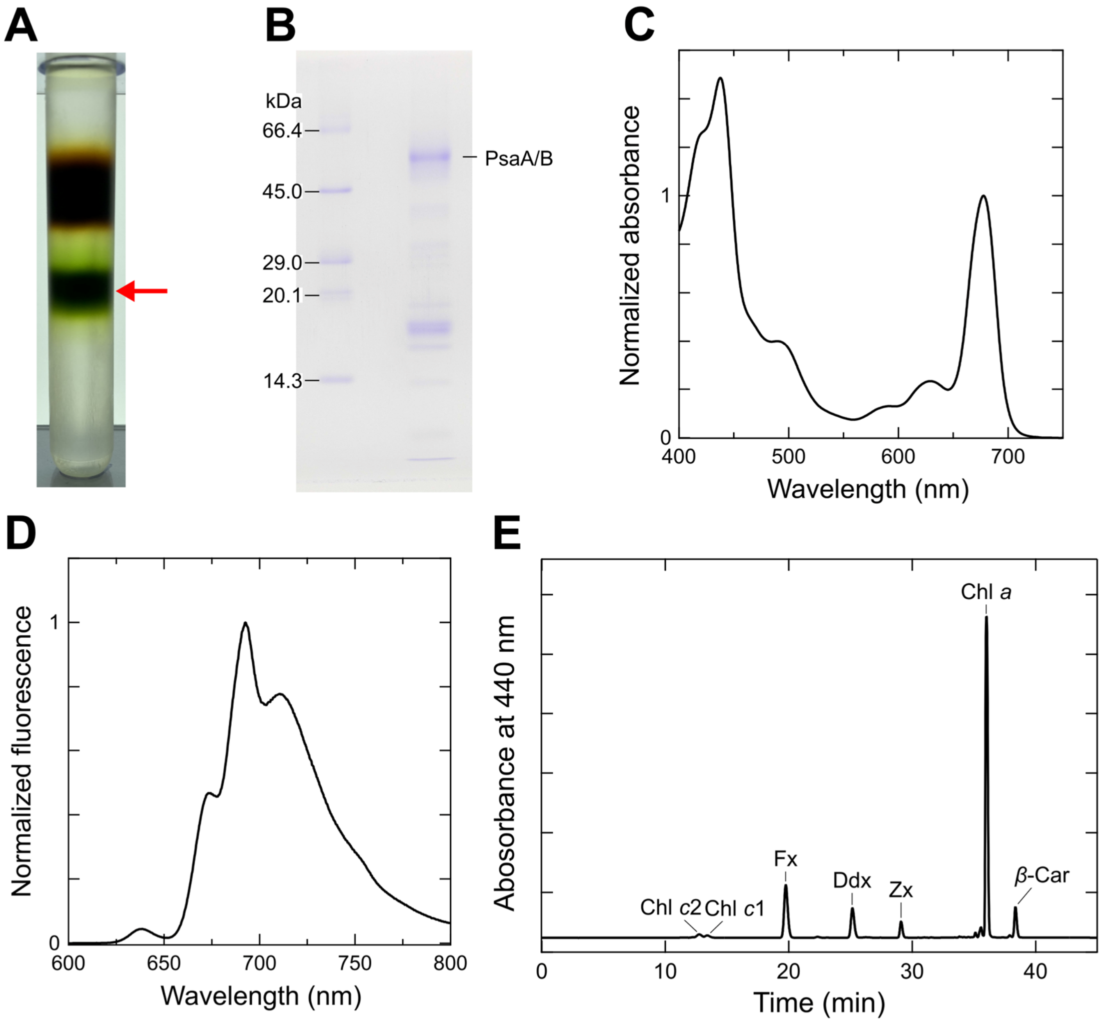


**Appendix 1—figure 1. Isolation and characterization of the *T. pseudonana* PSI-FCPI supercomplex.**

(**A**) Trehalose density gradient centrifugation. The red arrow indicates the PSI-FCPI fraction. (**B**) SDS-PAGE analysis of PSI-FCPI. PsaA/B proteins were tentatively identified by comparing their apparent molecular weights with markers. (**C**) Absorption spectrum of PSI-FCPI measured at room temperature. Three measurements were averaged, and the resulting spectrum was normalized by the intensity of the Qy peak. (**D**) Fluorescence-emission spectrum of PSI-FCPI measured at 77 K upon excitation at 430 nm. Three measurements were averaged, and the resulting spectrum was normalized by the maximum-peak intensity. (**E**) HPLC analysis of pigments extracted from PSI-FCPI, monitored at 440 nm. Chl *c*2, chlorophyll *c*2; Chl *c*1, chlorophyll *c*1; Fx, fucoxanthin; Ddx, diadinoxanthin; Zx, zeaxanthin; Chl *a*, chlorophyll *a*; *β*-Car, *β*-carotene. Data in panels **A**, **B**, and **E** are representative of three independent experiments.

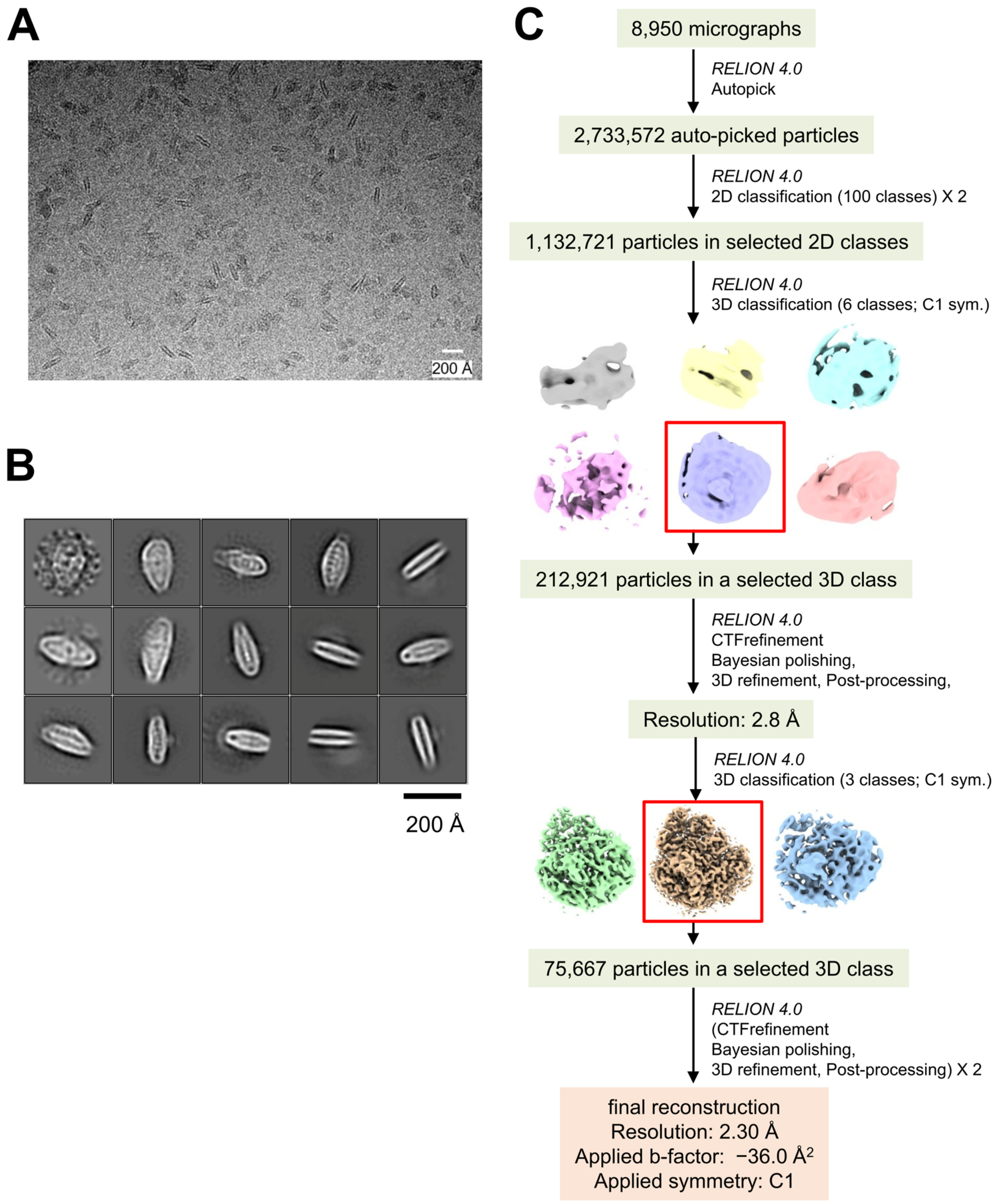


**Appendix 1—figure 2. Cryo-EM data collection and processing of PSI-FCPI.**
(**A**) A representative cryo-EM micrograph of PSI-FCPI from a total of 8,950 micrographs. (**B**) Representative 2D classes of PSI-FCPI. The box size is 300.8 Å. (**C**) Schematic flowchart illustrating the classification scheme and data processing for PSI-FCPI. Red boxes highlight selected particles from each 3D classification. The overall PSI-FCPI structure was reconstructed at a resolution of 2.30 Å from 75,667 particles. See the Methods section for further details.

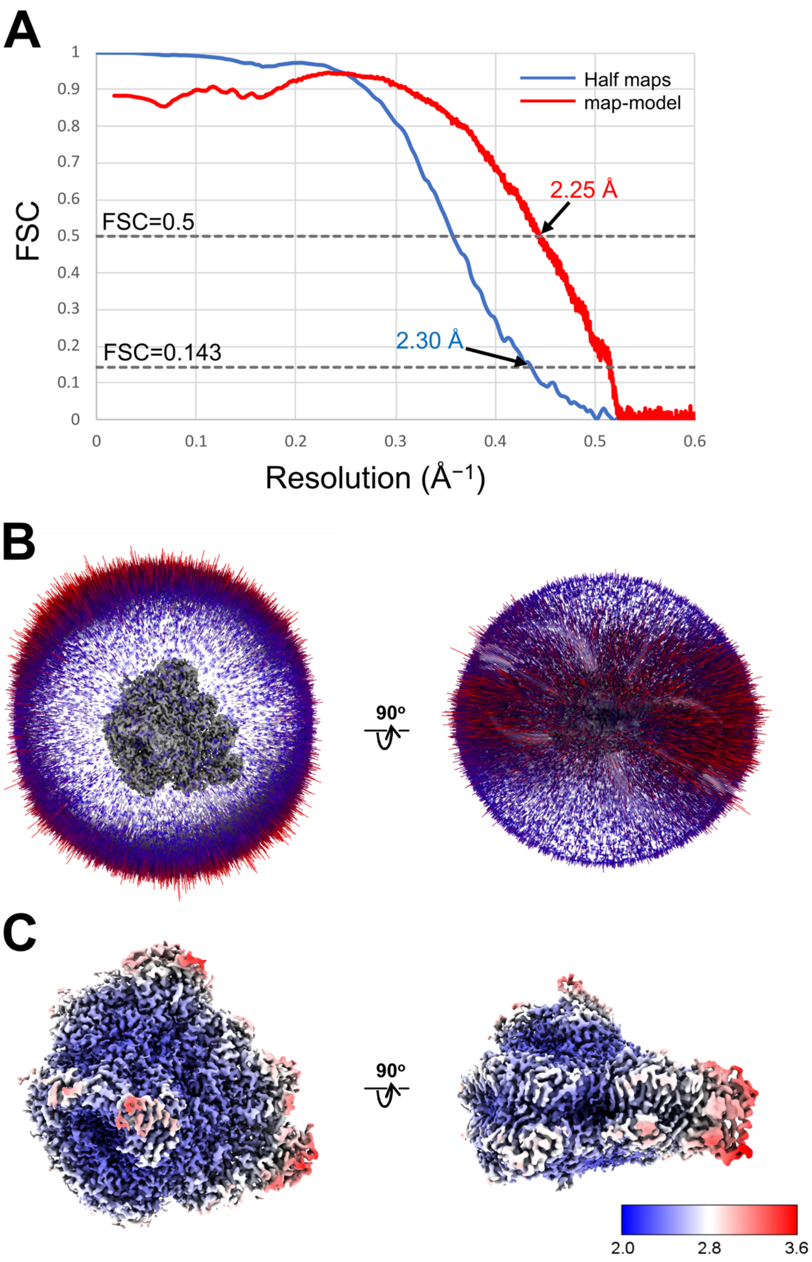


**Appendix 1—figure 3. Evaluation of the cryo-EM map quality.**
(**A**) FSC curves of PSI-FCPI for independently refined half maps (blue) and map-minus-model (red). (**B**) Angular distribution of the particles used for the reconstruction of PSI-FCPI. Each cylinder represents one view, and the height of the cylinder is proportional to the number of particles for that view. (**C**) Local resolution maps of PSI-FCPI.

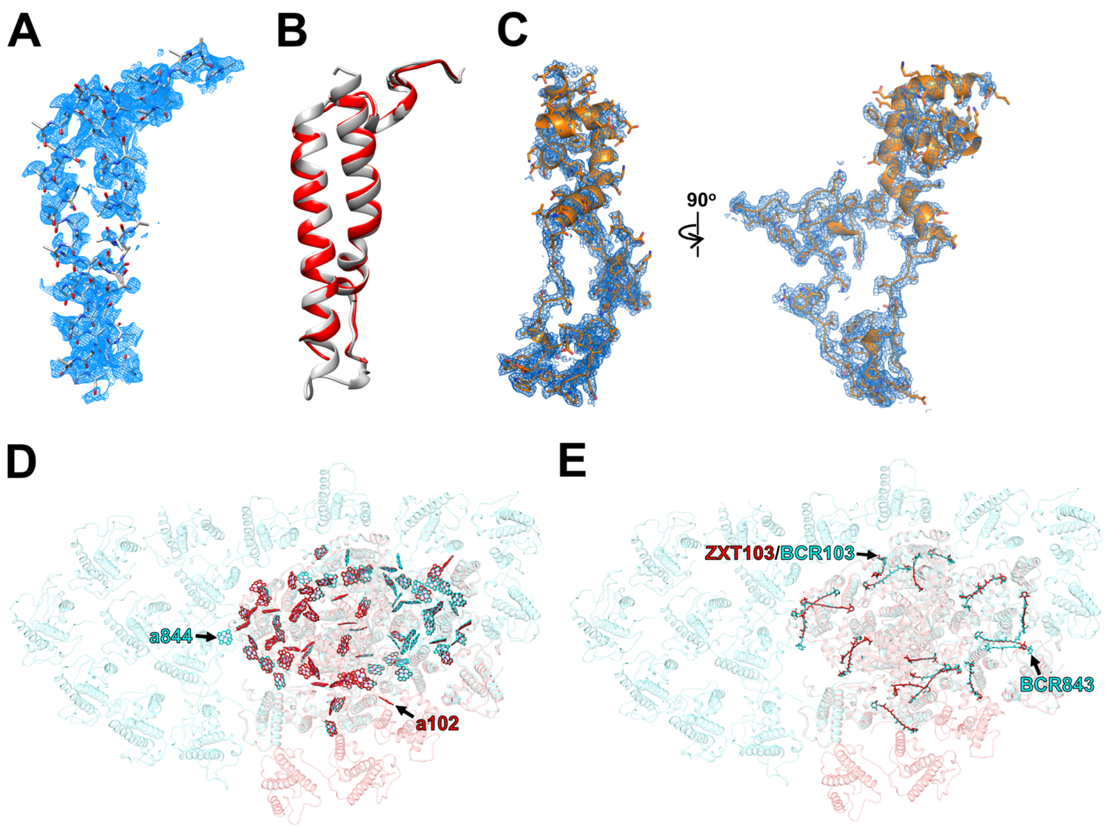


**Appendix 1—figure 4. Characteristic structures of the key PSI subunits and pigments in the PSI-FCPI structure.**

(**A**) The cryo-EM density for the Unknown subunit and its corresponding model are shown as meshes and sticks, respectively. (**B**) Superposition of protein structures between Unknown of *T. pseudonana* (red) and Psa28 of *C. gracilis* (grey) (PDB: 6L4U). (**C**) The cryo-EM density for Psa29 and its corresponding model are shown as meshes and sticks, respectively. (**D**, **E**) Comparison of Chls (**D**) and Cars (**E**) between *T. pseudonana* (red) and *C. gracilis* (cyan) PSI-FCPI structures. The *T. pseudonana* PSI-FCPI structure is superimposed on the *C. gracilis* PSI-FCPI structure (PDB: 6L4U), viewed from the stromal side. Chls and Cars are shown as sticks. Only rings of the Chl molecules are depicted. Characteristic pigments are labeled with red and cyan in the *T. pseudonana* and *C. gracilis* PSI-FCPI structures, respectively.

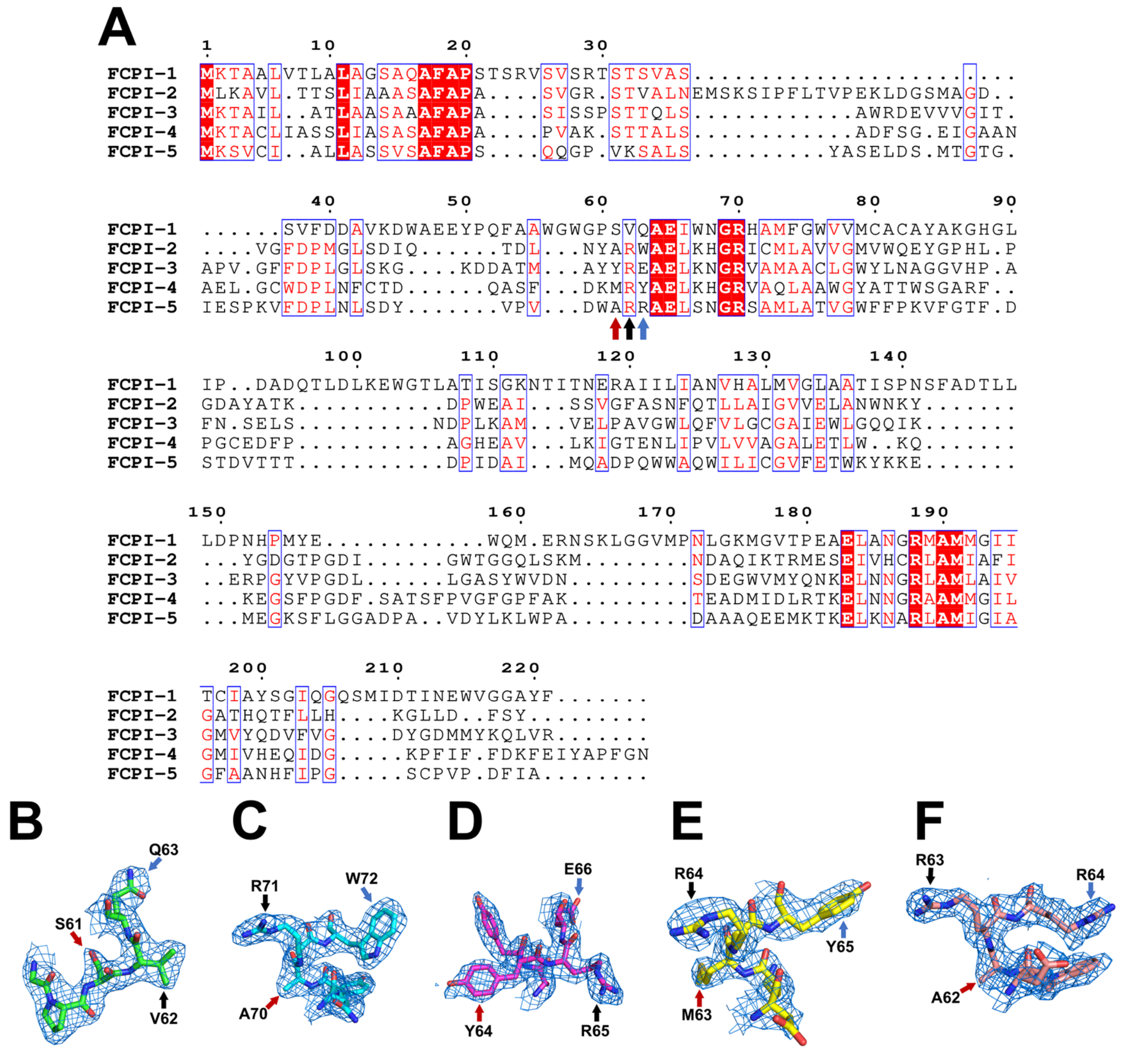


**Appendix 1—figure 5. Characteristic amino acid residues used for the identification of each FCPI subunit.**

(**A**) Multiple sequence alignment of FCPI proteins in *T. pseudonana* using PROMALS3D (http://prodata.swmed.edu/promals3d/promals3d.php) and ESPript (https://espript.ibcp.fr/ESPript/cgi-bin/ESPript.cgi). Unique residues are indicated by arrows in different colors, which were used to identify the various FCPI subunits. (**B–F**) Characteristic maps and amino acid residues of FCPI-1 (**B**), FCPI-2 (**C**), FCPI-3 (**D**), FCPI-4 (**E**), and FCPI-5 (**F**), respectively. The densities and models are shown as meshes and sticks, respectively. The characteristic amino acids are labeled with arrows in the same color as in panel **A**.

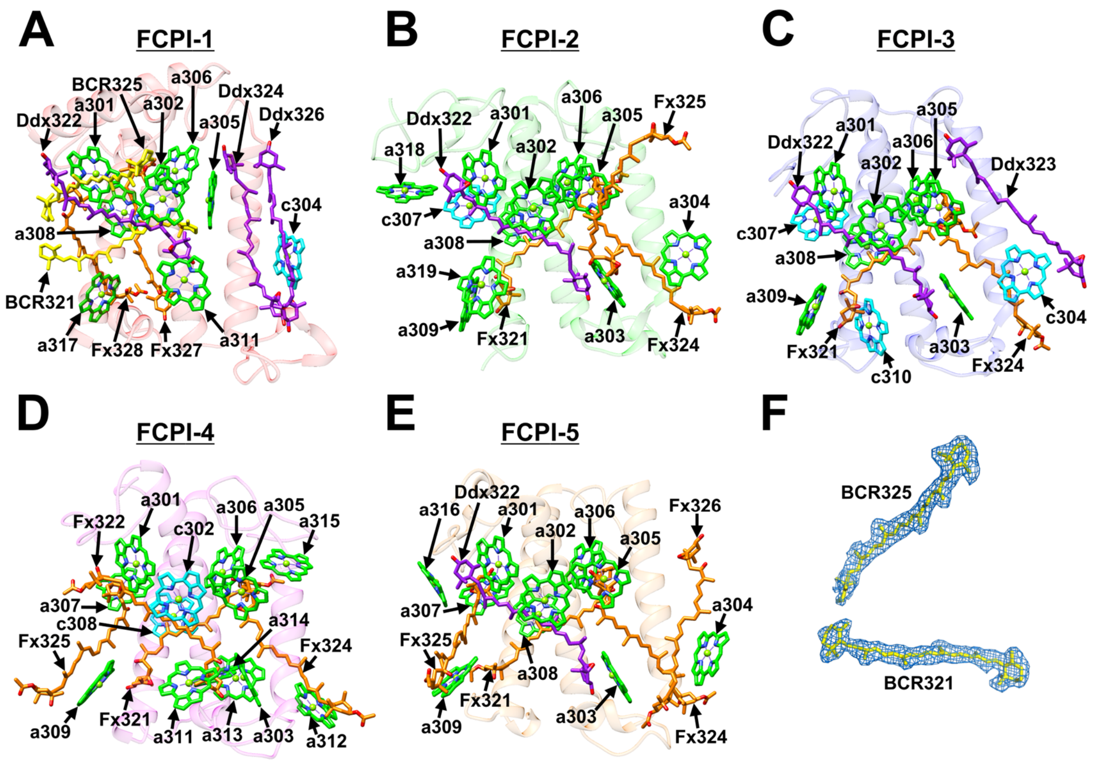

**Appendix 1—figure 6. Structures of FCPIs.**

(**A**–**E**) Structures of the five FCPI subunits from *T. pseudonana*, with proteins depicted as transparent cartoons and Chls and Cars shown as sticks in different colors. Only rings of the Chl molecules are depicted. Green, Chl *a*; cyan, Chl *c*; yellow, BCR; orange, Fx; purple, Ddx. (**F**) The cryo-EM densities for two BCRs in FCPI-1 and their corresponding models are shown as meshes and sticks, respectively.

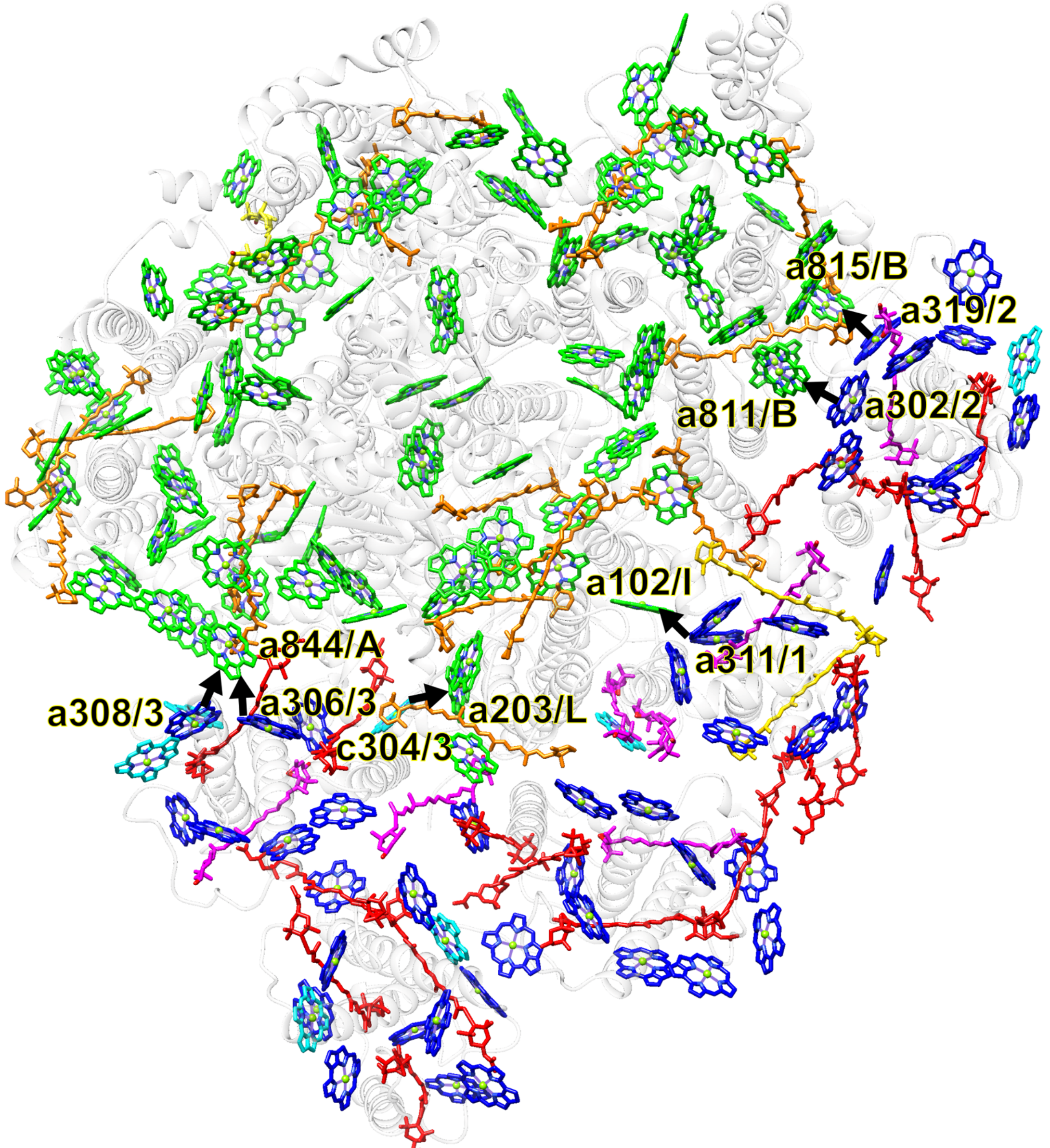


**Appendix 1—figure 7. Arrangement of pigment molecules within PSI-FCPI and possible excitation-energy-transfer pathways from FCPIs to PSI.**

The structure of the *T. pseudonana* PSI-FCPI is viewed from the stromal side. Chls and Cars are shown as sticks. Only rings of the Chl molecules are depicted. Black arrows indicate excitation-energy-transfer pathways based on close physical interactions among labeled Chls; for example, a844/A and c304/3 mean Chl a844 of PsaA and Chl c304 of FCPI-3, respectively. A, PsaA; B, PsaB; I, PsaI; L, PsaL; 1, FCPI-1; 2, FCPI-2; 3, FCPI-3. Green, Chls *a* in PSI; orange, BCRs in PSI; yellow, ZXT; blue, Chls *a* in FCPIs; cyan, Chls *c*; gold, BCRs in FCPI-1; red, Fxs; magenta, Ddxs.

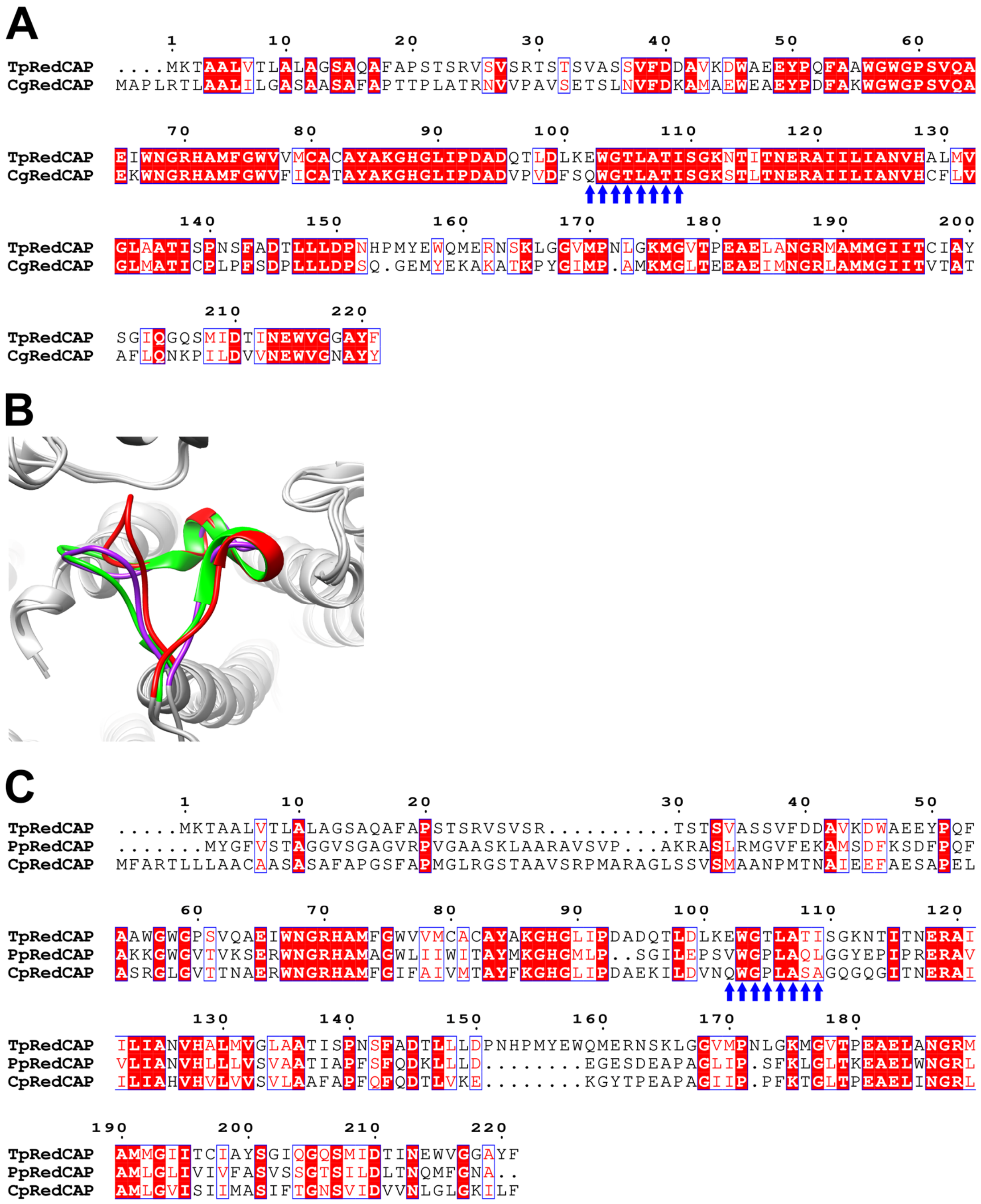


**Appendix 1—figure 8. Characteristics of the sequences and structures of RedCAPs in the red-lineage algae.**

(**A**) Sequence alignment of RedCAP of *T. pseudonana* (TpRedCAP) with that of *C. gracilis* CgRedCAP) using ClustalW (https://www.genome.jp/tools-bin/clustalw) and ESPript. Blue arrows indicate characteristic protein motifs (see text). (**B**) Structural comparisons of the Q96–T116 loop of TpRedCAP (red) with the corresponding loop of RedCAPs of *P. purpureum* (PpRedCAP; purple) (PDB: 7Y5E) and *C. placoidea* (CpRedCAP; green) (PDB: 7Y7B), viewed from the stromal side. (**C**) Multiple sequence alignment of TpRedCAP with PpRedCAP and CpRedCAP. Blue arrows indicate characteristic protein motifs (see text for more details).

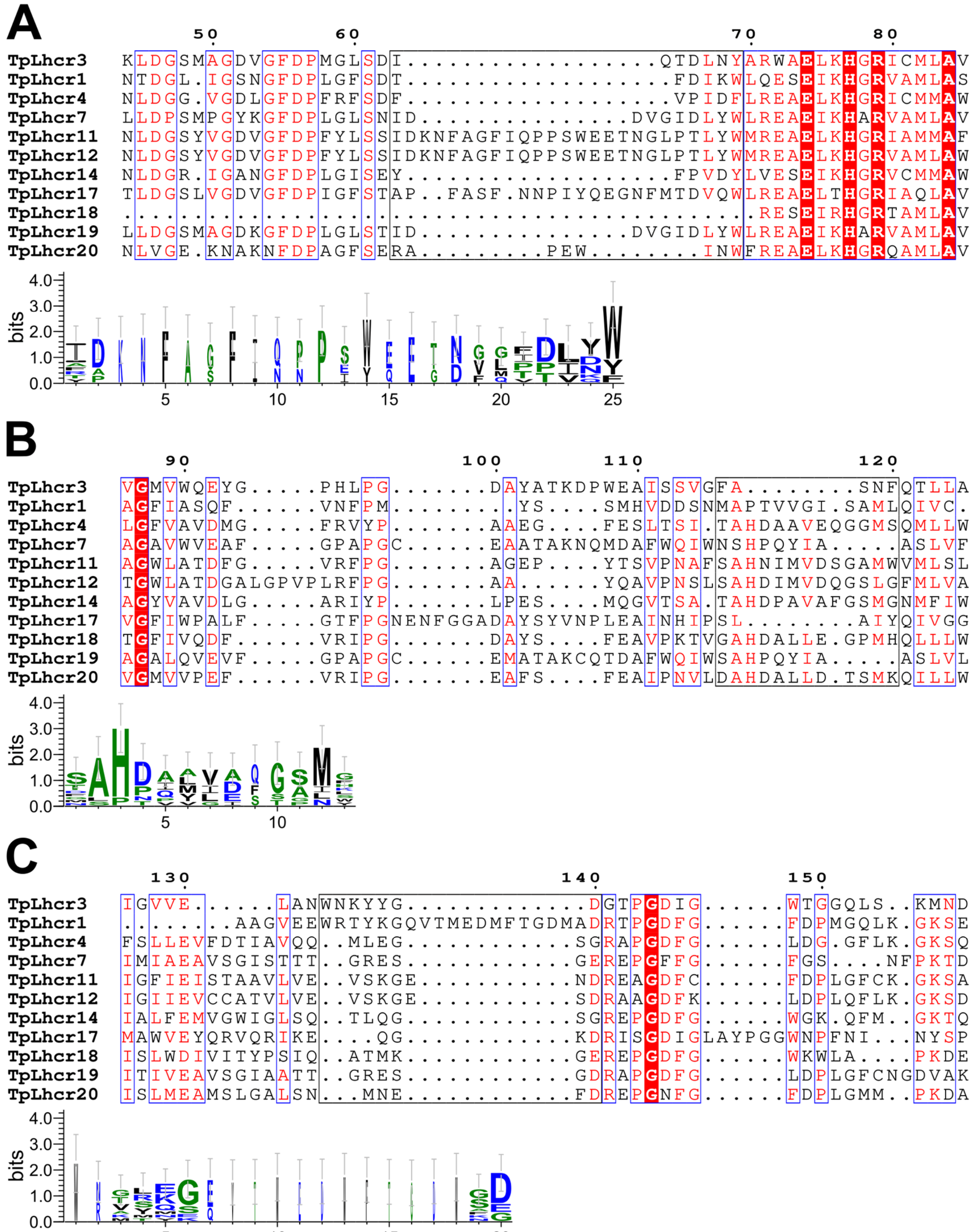


**Appendix 1—figure 9. Comparisons of the sequence of Lhcr3 (FCPI-2) with those of the Lhcr subfamily in *T. pseudonana*.**

Multiple sequence alignment of Lhcr3 of *T. pseudonana* (TpLhcr3) with the Lhcr subfamily (upper half of panels **A**–**C**). Amino acid residues I63–Y69 (**A**), F116–F120 (**B**), and W134–D140 (**C**) in Lhcr3, along with their corresponding residues in other Lhcrs, are highlighted with black boxes, and the consensus is displayed as sequence logos (lower half of each panel). Amino acid sequences were aligned using MAFFT E-INS-i v7.520 (https://mafft.cbrc.jp/alignment/software/). Sequence logos were generated by WebLogo v3.7.12 (https://weblogo.threeplusone.com/).

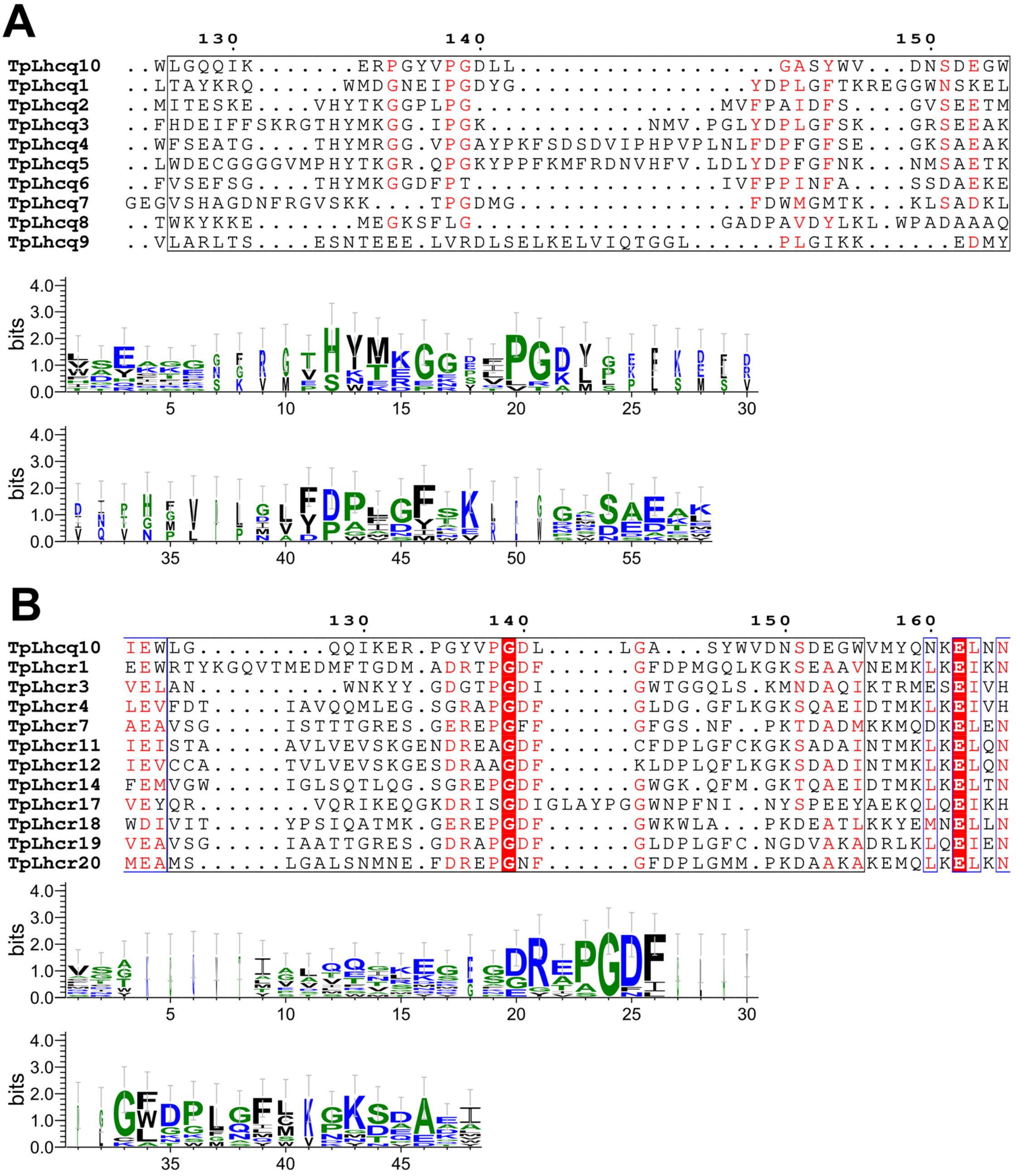


**Appendix 1—figure 10. Comparisons of the sequence of Lhcq10 (FCPI-3) with those of the Lhcq and Lhcr subfamilies in *T. pseudonana*.**

Multiple sequence alignments of Lhcq10 of *T. pseudonana* (TpLhcq10) with the Lhcq and Lhcr subfamilies (upper half of panels **A** and **B**, respectively). Amino acid residues L126–W155 in Lhcq10, along with their corresponding residues in other Lhcqs and Lhcrs, are highlighted with black boxes, and the consensus is displayed as sequence logos (lower half of each panel). Amino acid sequences were aligned using MAFFT E-INS-i v7.520. Sequence logos were generated by WebLogo v3.7.12.

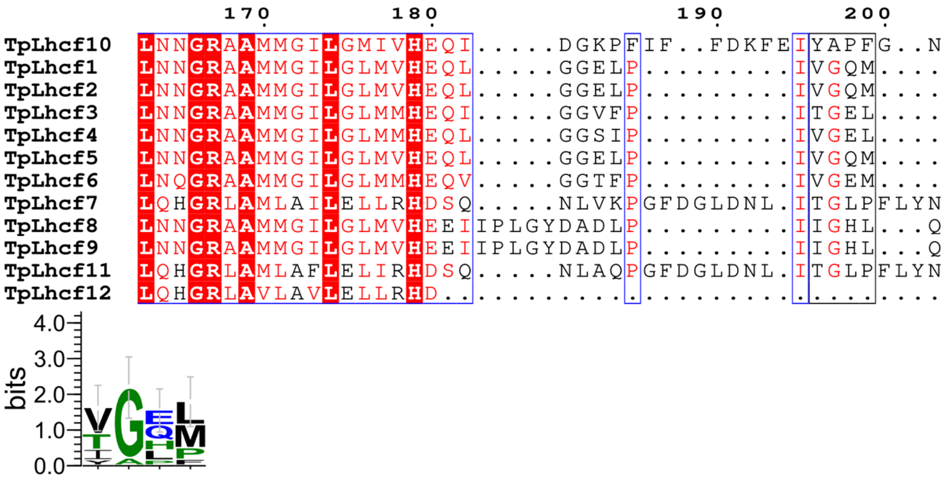

**Appendix 1—figure 11. Comparisons of the sequence of Lhcf10 (FCPI-4) with those of the Lhcf subfamily in *T. pseudonana*.**

Multiple sequence alignment of Lhcf10 of *T. pseudonana* (TpLhcf10) with the Lhcf subfamily (upper). Amino acid residues Y196–F199 in Lhcf10, along with their corresponding residues in other Lhcfs, are highlighted with a black box, and the consensus is displayed as sequence logos (lower). Amino acid sequences were aligned using MAFFT E-INS-i v7.520. Sequence logos were generated by WebLogo v3.7.12.

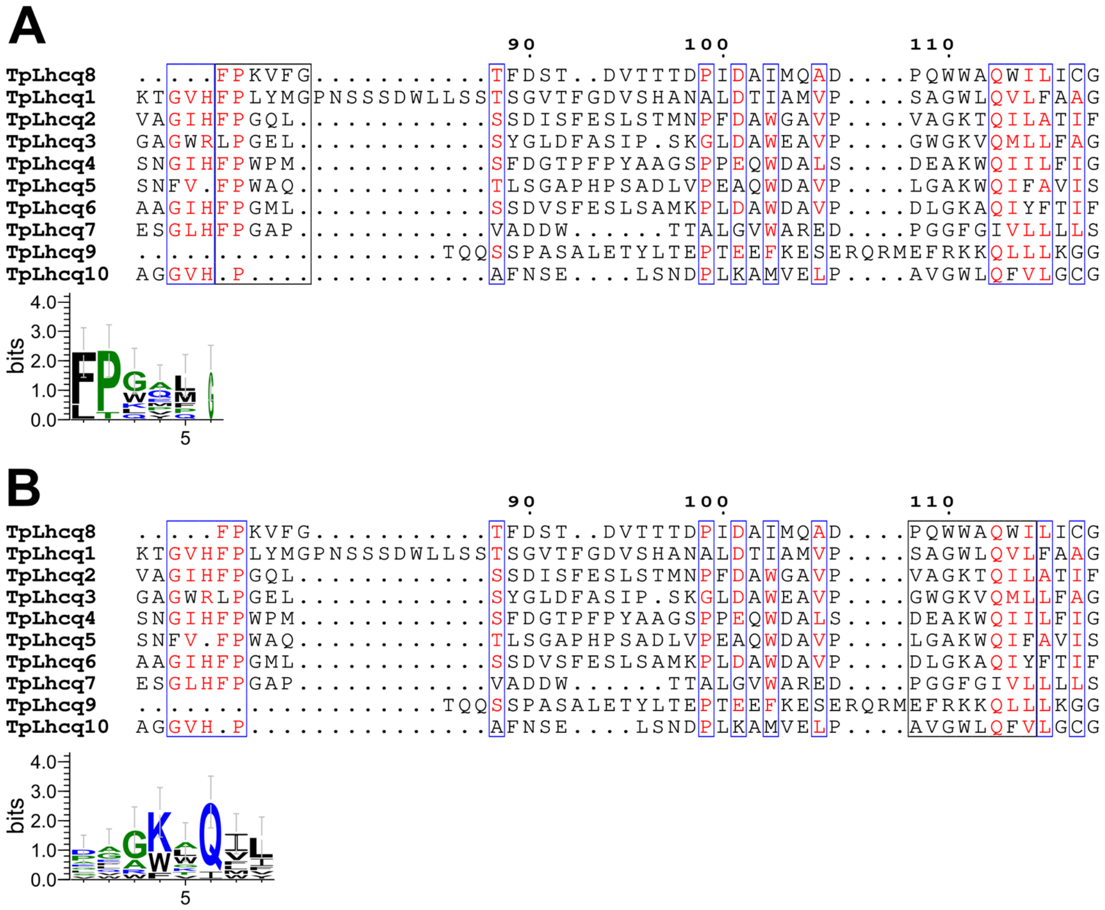

**Appendix 1—figure 12. Comparisons of the sequence of Lhcq8 (FCPI-5) with those of the Lhcq subfamily in *T. pseudonana*.**

Multiple sequence alignment of Lhcq8 of *T. pseudonana* (TpLhcq8) with the Lhcq subfamily (upper half of panels **A** and **B**). Amino acid residues F82–G87 and P107–I115 in Lhcq8, along with their corresponding residues in other Lhcqs, are highlighted with black boxes, and the consensus is displayed as sequence logos (lower half of each panel). Amino acid sequences were aligned using MAFFT E-INS-i v7.520. Sequence logos were generated by WebLogo v3.7.12.

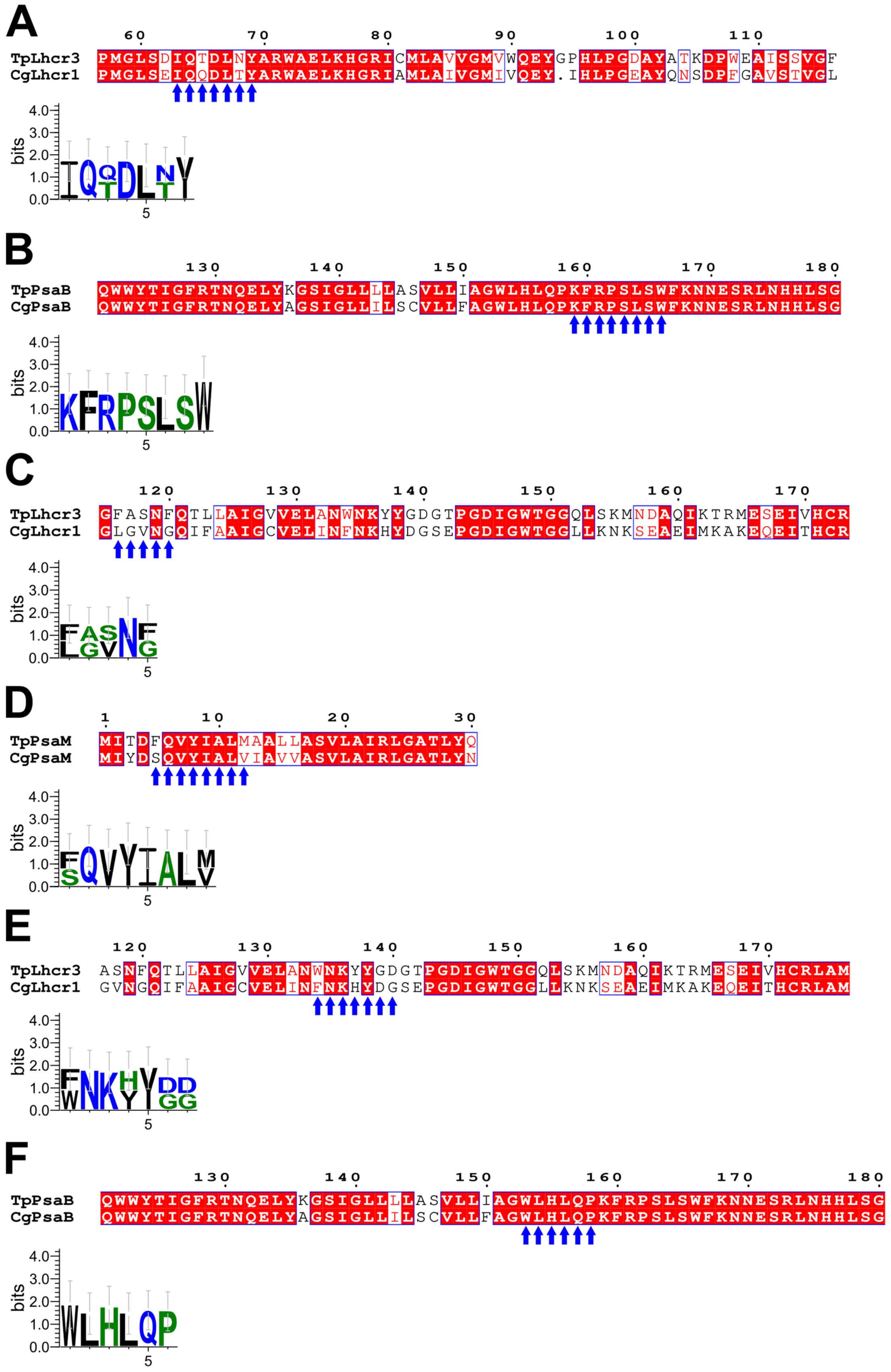


**Appendix 1—figure 13. Comparisons of amino acid sequences between FCPs and PSI proteins at the FCPI-2 site in the *T. pseudonana* PSI-FCPI structure.**

Sequence alignments between TpLhcr3 and Lhcr1 of *C. gracilis* (CgLhcr1) (upper half of panels **A**, **C**, and **E**), between TpPsaB and CgPsaB (upper half of panels **B** and **F**), and between TpPsaM and CgPsaM (upper half of panel **D**). Blue arrows indicate protein motifs involved in protein-protein interactions between TpLhcr3/CgLhcr1 and PsaB (**A**, **B**), between TpLhcr3/CgLhcr1 and PsaM (**C**, **D**), and between TpLhcr3/CgLhcr1 and PsaB (**E**, **F**), and the consensus is displayed as sequence logos (lower half of each panel). Amino acid sequences were aligned using MAFFT E-INS-i v7.520. Sequence logos were generated by WebLogo v3.7.12.

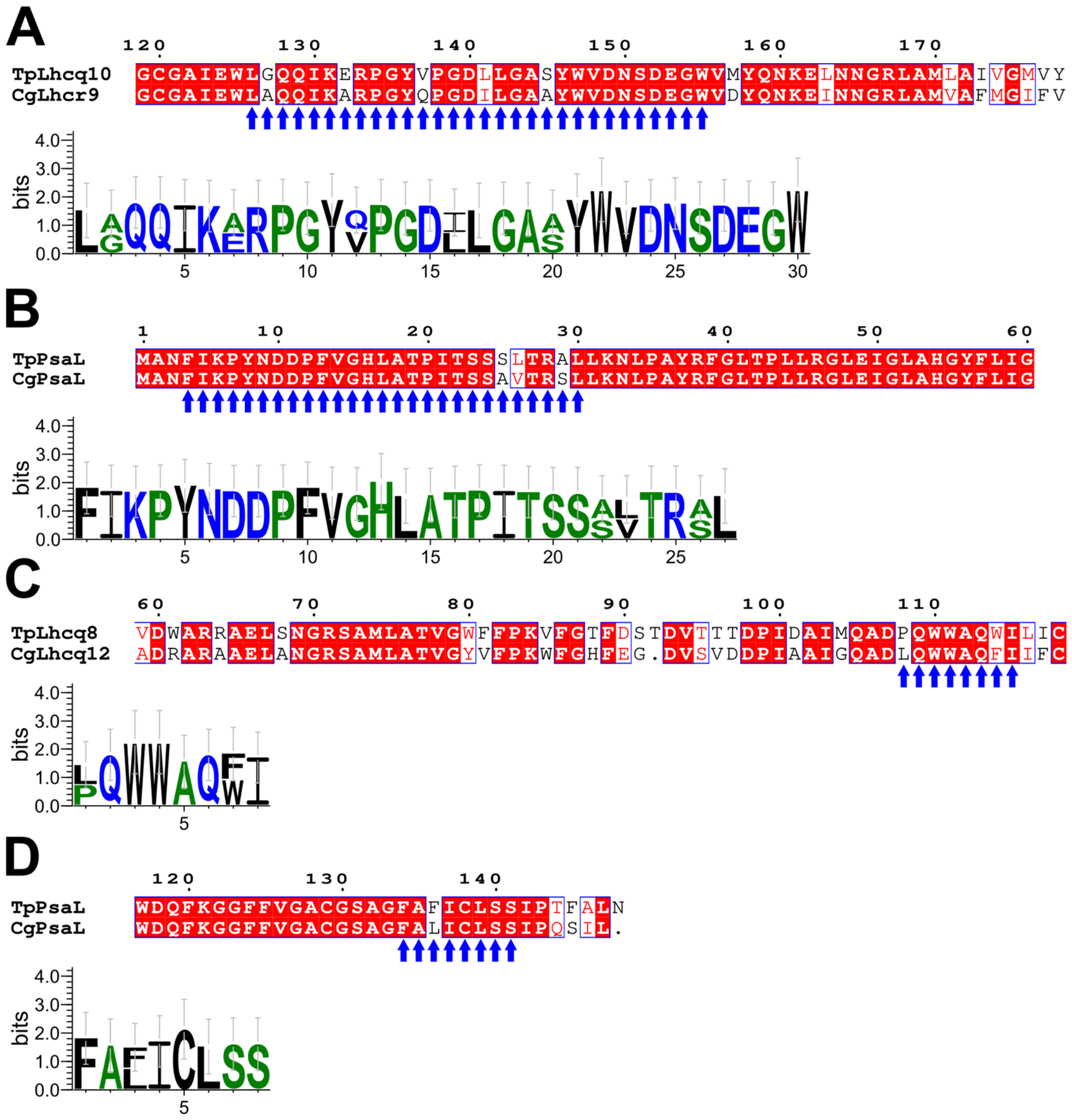


**Appendix 1—figure 14. Comparisons of amino acid sequences between FCPs and PsaL proteins at the FCPI-3 and FCPI-5 sites in the *T. pseudonana* PSI-FCPI structure.**

Sequence alignments between TpLhcq10 and CgLhcr9 (upper half of panel **A**), between TpPsaL and CgPsaL (upper half of panels **B**, **D**), and between TpLhcq8 and CgLhcq12 (upper half of panel **C**). Blue arrows indicate protein motifs involved in protein-protein interactions between TpLhcq10/CgLhcr9 and PsaL (**A**, **B**) and between TpLhcq8/CgLhcq12 and PsaL (**C**, **D**), and the consensus is displayed as sequence logos (lower half of each panel). Amino acid sequences were aligned using MAFFT E-INS-i v7.520. Sequence logos were generated by WebLogo v3.7.12.

**Appendix 1—table 1. Cryo-EM data collection and structural analysis statistics.**

| | |
|---|---|
| Complex | PSI-FCPI |
| PDB ID | 8XLS |
| EMDB ID | EMD-38457 |
| Data collection and processing | |
| Magnification | 60000 |
| Voltage (kV) | 300 |
| Electron exposure ($e^{-}$/Å) | 50 |
| Defocus range (µm) | −1.8 to −1.2 |
| Pixel size (Å) | 0.752 |
| Symmetry imposed | C1 |
| Initial particle images (no.) | 2,733,572 |
| Final particle images (no.) | 75,667 |
| Map resolution (Å) | 2.30 |
| FSC threshold | 0.143 |
| Refinement | |
| Initial Model used | De novo model building |
| Model resolution (Å) | 2.25 |
| FSC threshold | 0.5 |
| Map sharpening B factor ($Å^2$) | −36.0 |
| Model composition | |
| Non-hydrogen atoms | 37,640 |
| Protein residues | 3,129 |
| Ligand molecules | 372 |
| Water molecules | 922 |
| B factors ($Å^2$) | |
| Protein | 59.2 |
| Ligand | 71.9 |
| Water | 54.7 |
| R.m.s deviations | |
| Bond lengths (Å) | 0.025 |
| Bond angles (°) | 2.46 |
| Validation | |
| MolProbity score | 1.98 |
| Clashscore | 11.8 |
| Poor rotamers (%) | 3.04 |
| EMRinger score | 5.70 |
| Ramachandran plot | |
| Favored (%) | 97.90 |
| Allowed (%) | 2.07 |
| Disallowed (%) | 0.03 |

**Appendix 1—table 2. Averaged *Q*-scores in each subunit.**

| Subunit | Averaged *Q*-score | |
|---|---|---|
| | Postprocessed map | Denoised map |
| PsaA | 0.84 | 0.83 |
| PsaB | 0.84 | 0.83 |
| PsaC | 0.87 | 0.85 |
| PsaD | 0.83 | 0.83 |
| PsaE | 0.80 | 0.80 |
| PsaF | 0.81 | 0.81 |
| PsaI | 0.83 | 0.82 |
| PsaJ | 0.81 | 0.81 |
| PsaL | 0.83 | 0.83 |
| PsaM | 0.83 | 0.82 |
| Psa29 | 0.65 | 0.70 |
| Unknown | 0.45 | 0.55 |
| FCPI-1 | 0.76 | 0.78 |
| FCPI-2 | 0.67 | 0.72 |
| FCPI-3 | 0.74 | 0.76 |
| FCPI-4 | 0.77 | 0.79 |
| FCPI-5 | 0.74 | 0.76 |

**Appendix 1—table 3. Cofactors assigned in each subunit of the PSI-FCPI structure.**

| Protein | Chlorophyll | Carotenoid | Lipid | Other |
|---|---|---|---|---|
| PsaA | 43 Chl *a*<br>1 Chl *a′* | 5 BCR | 2 LHG | 1 [4Fe-4S] cluster<br>1 phylloquinone |
| PsaB | 41 Chl *a* | 5 BCR | 1 LHG<br>1 DGD | 1 phylloquinone |
| PsaC | - | - | - | 2 [4Fe-4S] cluster |
| PsaD | - | - | - | - |
| PsaE | - | - | - | - |
| PsaF | 3 Chl *a* | 1 BCR | - | - |
| PsaI | 1 Chl *a* | 1 BCR | - | - |
| PsaJ | 1 Chl *a* | 1 BCR<br>1 ZXT | - | - |
| PsaL | 3 Chl *a* | 3 BCR | 1 LMG | - |
| PsaM | - | 1 BCR | 1 LHG | - |
| Psa29 | - | - | - | - |
| Unknown | 1 Chl *a* | 1 BCR | - | - |
| FCPI-1 | 7 Chl *a*<br>1 Chl *c* | 2 BCR<br>2 Fx<br>3 Ddx | 1 LHG | - |
| FCPI-2 | 10 Chl *a*<br>1 Chl *c* | 3 Fx<br>1 Ddx | - | - |
| FCPI-3 | 7 Chl *a*<br>3 Chl *c* | 2 Fx<br>2 Ddx | 1 LHG | - |
| FCPI-4 | 11 Chl *a*<br>2 Chl *c* | 4 Fx | 1 LHG | - |
| FCPI-5 | 10 Chl *a* | 4 Fx<br>1 Ddx | - | - |
| Total | 146 | 43 | 9 | 5 |

BCR, *β*-carotene; ZXT, zeaxanthin; Fx, fucoxanthin; Ddx, diadinoxanthin; Chl *a*, chlorophyll *a*; Chl *a′*, chlorophyll *a* epimer; Chl *c*, chlorophyll *c*; DGD, digalactosyl diacyl glycerol; LHG, dipalmitoyl phosphatidyl glycerol; LMG, distearoyl monogalactosyl diglyceride.

**Appendix 1—table 4. FCPI proteins identified in the PSI-FCPI structure, their corresponding genes, and their RMSD values compared with the FCPI-4 structure.**

| Protein | Gene | RMSD (Å)/Aligned Cα atoms |
|---|---|---|
| FCPI-1 | *RedCAP* | 3.73/95 |
| FCPI-2 | *Lhcr3* | 2.01/139 |
| FCPI-3 | *Lhcq10* | 2.02/139 |
| FCPI-4 | *Lhcf10* | 0.00/167 |
| FCPI-5 | *Lhcq8* | 1.91/128 |

**Appendix 1—table 5. Chls and their ligands in each of the FCPI subunits.**

| Protein | Chlorophyll/ligand |
|---|---|
| FCPI-1 | a301/E65, a302/N68, c304/H128, a305/H71, a306/E183, a308/N186, a311/w982[2], a317/W215 |
| FCPI-2 | a301/E74, a302/H77, a303/Q91, a304/Q121, a305/E130, a306/E168, c307/-[1], a308/H171, a309/Q185, a318/S35, a319/H184 |
| FCPI-3 | a301/E68, a302/N71, a303/w977[2], c304/Q115, a305/E124, a306/E162, c307/w976[2], a308/N165, a309/w978[2], c310/D188 |
| FCPI-4 | a301/E67, c302/H70, a303/-[1], a305/E122, a306/E162, a307/LHG330, c308/N165, a309/H179, a311/H100, a312/P91, a313/w980[2], a314/Y196, a315/P142 |
| FCPI-5 | a301/E66, a302/N69, a303/w994[2], a304/Q113, a305/E122, a306/E163, a307/E43, a308/N166, a309/S186, a316/E43 |

[1]The ligands of Chls may be water or lipid molecules which cannot be identified due to weak densities.

[2]Water molecules.

**Appendix 1—table 6. Correspondence of the numbering of pigments in each PSI core subunit described in the text with those in the PDB file.**

| | PsaI | PsaJ | PsaL |
|---|---|---|---|
| **Chls in the text** | **PDB No. (Chain ID)** | **PDB No. (Chain ID)** | **PDB No. (Chain ID)** |
| 102 | 301 (1)* | | |
| 203 | | | 204 (L) |
| **Car in the text** | | | |
| 103 | | 105 (J) | |

*Chain in the adjacent unit.

**Appendix 1—table 7. Correspondence of the numbering of pigments in each FCPI subunit described in the text with those in the PDB file.**

| | FCPI-1 | FCPI-2 | FCPI-3 | FCPI-4 | FCPI-5 |
|---|---|---|---|---|---|
| **Chls in the text** | **PDB No. (Chain ID)** | **PDB No. (Chain ID)** | **PDB No. (Chain ID)** | **PDB No. (Chain ID)** | **PDB No. (Chain ID)** |
| 301 | 303 (1) | 205 (2) | 202 (3) | | 207 (5) |
| 302 | 304 (1) | 206 (2) | 203 (3) | | 208 (5) |
| 303 | | 207 (2) | 204 (3) | | 209 (5) |
| 304 | 305 (1) | 208 (2) | 205 (3) | | 210 (5) |
| 305 | 306 (1) | 209 (2) | 206 (3) | 304 (4) | 211 (5) |
| 306 | 307 (1) | 210 (2) | 207 (3) | 305 (4) | 212 (5) |
| 307 | | 211 (2) | 208 (3) | 306 (4) | 213 (5) |
| 308 | | 212 (2) | 209 (3) | 307 (4) | 214 (5) |
| 309 | | 213 (2) | 210 (3) | 308 (4) | 215 (5) |
| 310 | | | 211 (3) | | |
| 311 | 309 (1) | | | 309 (4) | |
| 312 | | | | 310 (4) | |
| 313 | | | | 311 (4) | |
| 314 | | | | 312 (4) | |
| 315 | | | | 313 (4) | |
| 316 | | | | | 216 (5) |
| 317 | 310 (1) | | | | |
| 318 | | 214 (2) | | | |
| 319 | | 215 (2) | | | |
| **Cars in the text** | | | | | |
| 321 | 311 (1) | 216 (2) | 212 (3) | 314 (4) | 217 (5) |
| 322 | 312 (1) | 217 (2) | 213 (3) | 315 (4) | 218 (5) |
| 323 | | | 209 (L)* | | |
| 324 | 313 (1) | 218 (2) | 214 (3) | 316 (4) | 219 (5) |
| 325 | 314 (1) | 219 (2) | | 228 (3)* | 220 (5) |
| 326 | 315 (1) | | | | 221 (5) |
| 327 | 316 (1) | | | | |
| 328 | 317 (1) | | | | |

*Chain in the adjacent unit.

**Appendix 1—table 8. Correspondence of the numbering of other cofactors described in the text with those in the PDB file.**

| Waters in the text | PDB No. (Chain ID) |
|---|---|
| 976 | 317 (3) |
| 977 | 313 (3) |
| 978 | 316 (3) |
| 980 | 402 (4) |
| 982 | 425 (1) |
| 994 | 308 (5) |

| Lipid in the text | |
|---|---|
| 330 | 317 (4) |
